# Structural basis for the specific recognition of DSR by the YTH domain containing protein Mmi1

**DOI:** 10.1101/145276

**Authors:** Baixing Wu, Jinhao Xu, Shichen Su, Hehua Liu, Jianhua Gan, Jinbiao Ma

**Author notes:** Co-first authors (Baixing Wu, Jinhao Xu).

## Abstract

Meiosis is one of the most dramatic differentiation programs accompanied by a striking change in gene expression profiles, whereas a number of meiosis-specific transcripts are expressed untimely in mitotic cells. The entry of meiosis will be blocked as the accumulation of meiosis-specific mRNAs during the mitotic cell in fission yeast *Schizosaccharomyces pombe*. A YTH domain containing protein Mmi1 was identified as a pivotal effector in a post-transcriptional event termed selective elimination of meiosis-specific mRNAs, Mmi1 can recognize and bind a class of meiosis-specific transcripts expressed inappropriately in mitotic cells, which contain a conservative motif called DSR as a mark to remove them in cooperation with nuclear exosomes. Here we report the 1.6 Å resolution crystal structure of the YTH domain of Mmi1 binds to high-affinity RNA targets r(A_1_U_2_U_3_A_4_A_5_A_6_C_7_A_8_) containing DSR core motif. Our structure observations, supported by site-directed mutations of key residues illustrate the mechanism for specific recognition of DSR-RNA by Mmi1. Moreover, different from other YTH domain family proteins, Mmi1 YTH domain has a distinctive function although it has a similar fold as other ones.

## Introduction

Meiosis is an important cellular differentiation process in which a group of transcripts are newly induced or down-regulated. Amounts of meiosis-specific transcripts that are abundant in meiosis can be detected in early mitosis but be quickly removed(Harigaya et al. 2006). Meiotic-specific genes must be tightly regulated in the mitotic cell cycle because of the expression of meiotic genes during mitosis is detrimental to proliferation. A protein named Mmi1 plays important role in the post-transcriptional event termed selective elimination of meiosis-specific mRNAs. In mitosis, Mmi1 recognizes a number of meiosis-specific transcripts that are expressed untimely in mitotic cells and removes them in cooperation with nuclear exosomes. These transcripts carry a region called DSR (Determinant of Selective Removal), to which Mmi1 binds. Cells would not continue robust mitotic proliferation due to the accumulation of deleterious meiosis-specific transcripts if this elimination system has something wrong within it. In meiosis, Mmi1 inhibits the progression of meiosis because Mmi1 regulates meiotic transcripts that are essential for meiosis, moreover, Mmi1 overexpression impairs meiosis(Chen et al. 2011). To overcome Mmi1 mediated suppression of meiosis, another RNA-binding protein, Mei2, and its binding partner, meiRNA, suppress Mmi1. In meiotic prophase, Mei2 and meiRNA form a dot structure at the sme2 gene locus on chromosome II, which encodes meiRNA. MeiRNA carries multiple copies of the DSR motif and is degraded via the Mmi1 mediated degradation machinery(Harigaya et al. 2006; Shichino et al. 2014). On the basis of before observations, meiRNA may serve as a decoy substrate for Mmi1 (Shichino et al. 2014). Yet, little is known about mechanisms used by mitotic cells to repress meiosis-specific genes. The recognition of Mmi1 and DSR motif comes to the key point to demystify the mechanism.

Mmi1 consists of a C-terminal YTH domain, which specifically binds to DSR sequence (Yamashita et al. 2012). RNA binding proteins of the YTH family appear to be conserved widely among eukaryotes, prompting an important function of the YTH domain across species. Rat YT521-B, the founding member of this family, has been shown to interact with several splicing factors and to alter alternative splice sites in a dose-dependent manner(Stoilov et al. 2002). Recent studies showed that YTH domain family protein 2 (YTHDF2) and other YTH domain containing protein, *e.g.* MRB1 in budding yeast and *Hs*YTHDC1, preferentially bind to m^6^A-containing mRNA *in vivo* and *in vitro* and regulate localization and stability of the bound mRNA(Schwartz et al. 2013; Dominissini et al. 2012; Wang et al. 2014). The YTH domain contains about 160 residues usually located at the C-terminal, and the N-terminal regions of these proteins are poorly conserved.

Yamashita *et al*. identify the nucleotide motifs that are essential for DSR function using both a computational method known as “motif sampling analysis” and a genetic method involving mutational analysis(Chen et al. 2011). They show that Mmi1 interacts with repeats of the hexanucleotide 5’-U(U/C)AAAC-3’ that are enriched in the DSR. Disruption of this “DSR core motif” in a target mRNA inhibits its elimination. Tandem repeats of the motif can function as an artificial DSR, Mmi1 binds to it *in vitro*. A core motif cluster may responsible for the DSR activity. In order to explore selective elimination system by Mmi1-DSR more profoundly and if Mmi1 has the same function to recognize the m^6^A modification, we expressed and purified the Mmi1 YTH domain *in vitro*, and further crystallized the Mmi1 in complex with DSR core motif 5’-A(UUAAACA)A-3’ to illuminate the detail mechanism for recognition of RNA targets by Mmi1.

## RESULTS

### Crystal structure of Mmi1 YTH domain

Similar to most YTH domain family proteins in other species, the conserved YTH domain of Mmi1 from *S. pombe* is located in its C-terminal region (Figure 1A). Based on the sequence alignment (Figure 1B), the construct of Mmi1 (amino acids 349–477) containing most conserved part has been chosen to be expressed in *E. coli*, purified to homogeneous through Ni-chelating and gel-filtration columns, and successfully crystallized after screening. The structure has been solved by single-wavelength anomalous dispersion (SAD) method using selenomethionine-labeled crystal diffracted to 2.0 Å resolution and refined to 1.49 Å resolution using diffraction data from a native crystal (Table 1). Because this shorter construct of Mmi1 YTH domain failed to bind DSR-RNA (data not shown), the crystal structure of longer construct (amino acids 326–488) that binds DSR-RNA very well (Figure 2A), was determined at 1.8 Å resolution by molecular replacement (MR) method (Table 1).

**Figure 1.**
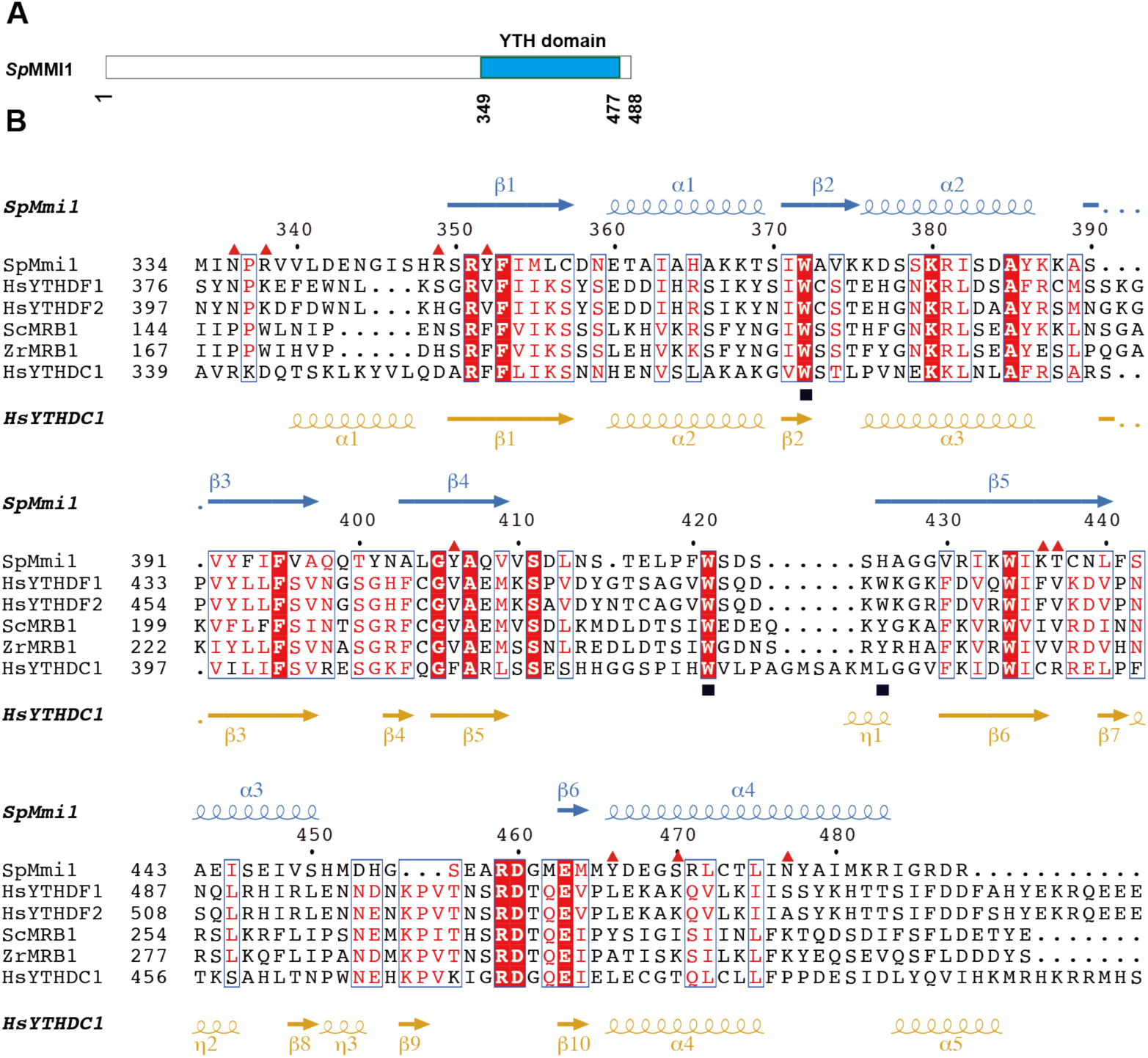
Domain architecture and sequence alignment. (A) Domain architecture of the Mmi1 construct (349 to 477) consisting of the YTH domain (in blue) for crystallization of the complex. (B) Sequence alignment of the YTH domain family members. Sequence alignment between YTH domain of *S. pombe* Mmi1, *S. cerevisiae* MRB1, and *H. sapiens* YTHDC1, YTHDF1 and YTHDF2. Secondary structure alignments of Mmi1 and YTHDC1 YTH domain is shown above (in blue) and below (in yellow) the sequences, respectively. Numbering above the sequences corresponds to the *S. pombe* Mmi1. Red triangles denote residues that form hydrogen bonds with RNA bases or sugar phosphate backbone. Black rectangles indicate the residues known as m6A binding pocket important to HsYTHDC1 for recognizing the m^6^A target RNA.

**Figure 2.**
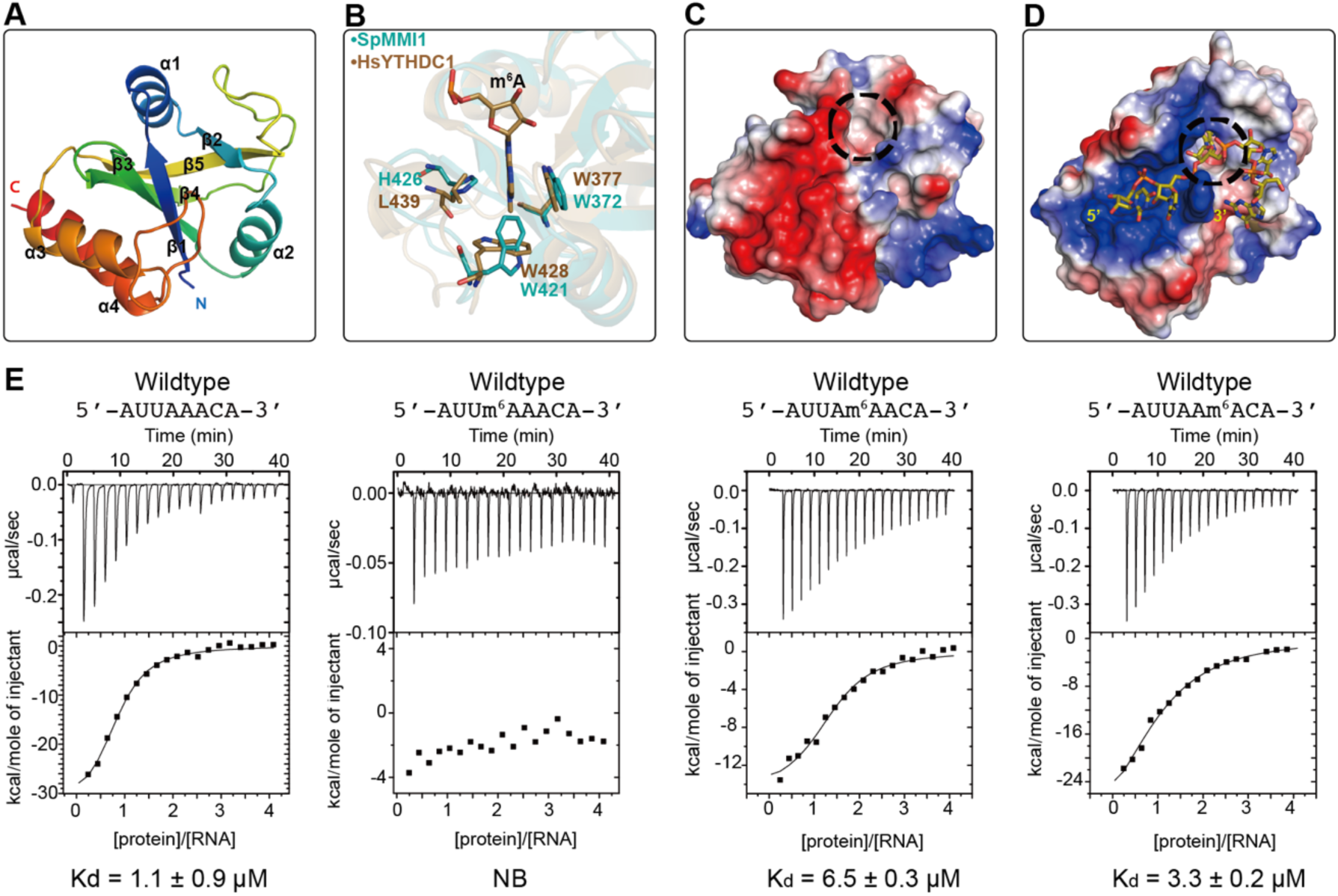
Mmi1 YTH domain does not recognize m^6^A RNA. **(A)** Apo structure of Mmi1-YTH domain (349–477) are shown as cartoon in rainbow. (B) Superposition of the m^6^A binding pocket of YTHDC-YTH domain with Mmi1-YTH domain. The aromatic cage residues and m6A RNA of YTHDC-YTH domain are shown in brown sticks, the correspondence residues of the Mmi1-YTH domain are shown in cyan sticks. (C-D) Electrostatic surface potential comparison of YTHDC1-YTH-RNA complex and Mmi1-YTH at the same perspective. The aromatic cage is circled with black dashed lines. (E) ITC binding curves show the Mmi1-YTH domain preference bind to DSR core motif 5’-AUUAAACA-3’ but not m^6^A modified RNA targets. N^6^ methylated target RNA sequence as follows: 5-AUU(m^6^A)AACA-3’, 5-AUUA(m^6^A)ACA-3’, 5-AUUAA(m^6^A)CA-3’.The upper panels show the raw calorimetric data. The bottom plots are integrated heats as a function of the YTH/RNA molar ration. Black dots indicate the experimental data. The best fit (depicted by the black line) was obtained from a non-linear least-squares method using a one-site binding model.

The overall structure of the Mmi1 YTH domain adopts a globular fold with a central core of five-stranded β-sheets surrounded by four α-helices and flanking regions (Figure 2A). The root mean square deviation (RMSD) of the backbone Cα atoms between Mmi1 YTH domain and other YTH domains are small, e.g. 1.068 Å with YTHDC1, 1.311 Å with *Zr*MRB1, 1.592 Å with YTHDF1, suggesting that the Mmi1 YTH domain maintain the overall structural fold similar to reported YTH domains (Figure S1). In addition, Mmi1 YTH domain contains a potential m^6^A binding pocket consisting of three aromatic residues, W372, W421 and H426, that are conserved in other reported YTH family proteins (Figure 1B and Figure 2B). However, the electrostatic potential surface of the Mmi1 YTH domain shows a large negatively charged patch surrounding the potential m^6^A binding pocket (Figure 2C), whereas a strong positively charged patch exhibits on the surfaces of YTH domains binding m^6^A RNA (Figure 2C, 2D and Figures S2A, S2B and S2C).

Because all the YTH domains have reported binding m^6^A modified RNA, we measured the binding affinities between Mmi1 YTH domain with six different m^6^A modified RNA containing GG(m^6^A)CU or DSR core motif carried m^6^A modification. However, no binding can be detected by EMSA or ITC (data not shown). The H426 and W372 in Mmi1 YTH domain have the same conformation with corresponding tryptophan in other YTH domains. The most significant difference between Mmi1 and the other three YTH domains is W421, of which the indole ring turns almost 90 degrees relative to the other three protein’s tryptophan (Figure 2B). In addition, the electrostatic surface is the critical reason determines Mmi1 YTH domain has no ability to recognize the GG(m6A)CU motif. Our ITC results show that the Mmi1 YTH domain binds DSR with high binding affinity, although the binding affinities decrease one by one while the m^6^A modification turns from A6 to A4 (Figure 2E). These results suggest that Mmi1 YTH domain does not recognize the m^6^A modification in DSR. The A1 and A8 is not contained in the core DSR motif, though we cannot get crystals without these two residues,

### Crystal structure of Mmi1 YTH domain-DSR complex

We then obtained the crystal structure of Mmi1-YTH domain in complex with DSR containing RNA without m^6^A modification. The crystal belongs to the space group of P2_1_2_1_2_1_ and the structure is determined by molecular replacement and refined to 1.6 Å resolution (Table 1). We named the RNA sequence from 5’-3’ as A1 to A8 (Figure 3A). The conformation has not dramatically change to the apo-Mmi1 structure except an extended loop flanking at the N-terminus, and also the aromatic cage corresponding to the m^6^A binding is keep unchanged, although there is one residue different from the other YTH domain proteins, A362, which can recognize the N^1^ of the m^6^A (Figure S3A, S3B). The RNA substrate lies in a groove formed by α4, β3, and β4 full of positive charge(Figure 3A, 3B). The RNA backbone adopts a turn between U3 and A4 (Figure 3C), the base of A4 stacks over a hydrophobic pocket formed by Y392, Y406 and C473, and the bases of A4 to A8 continuously stack on each other along the groove surface (Figure 3D). The residue K437 forms the only interaction with the backbone phosphate of the RNA (Figure 3E and 3F). Consistent with the consensus motif 5‘-U(U/C)AAAC-3’ identified by microarray from earlier studies, there is no interaction between U3 and the Mm1 YTH domain, though U3 is stacked by A1. A4 occupies the central position, suggesting that the A4 nucleotide is pivotal for protein-RNA recognition.

**Figure 3.**
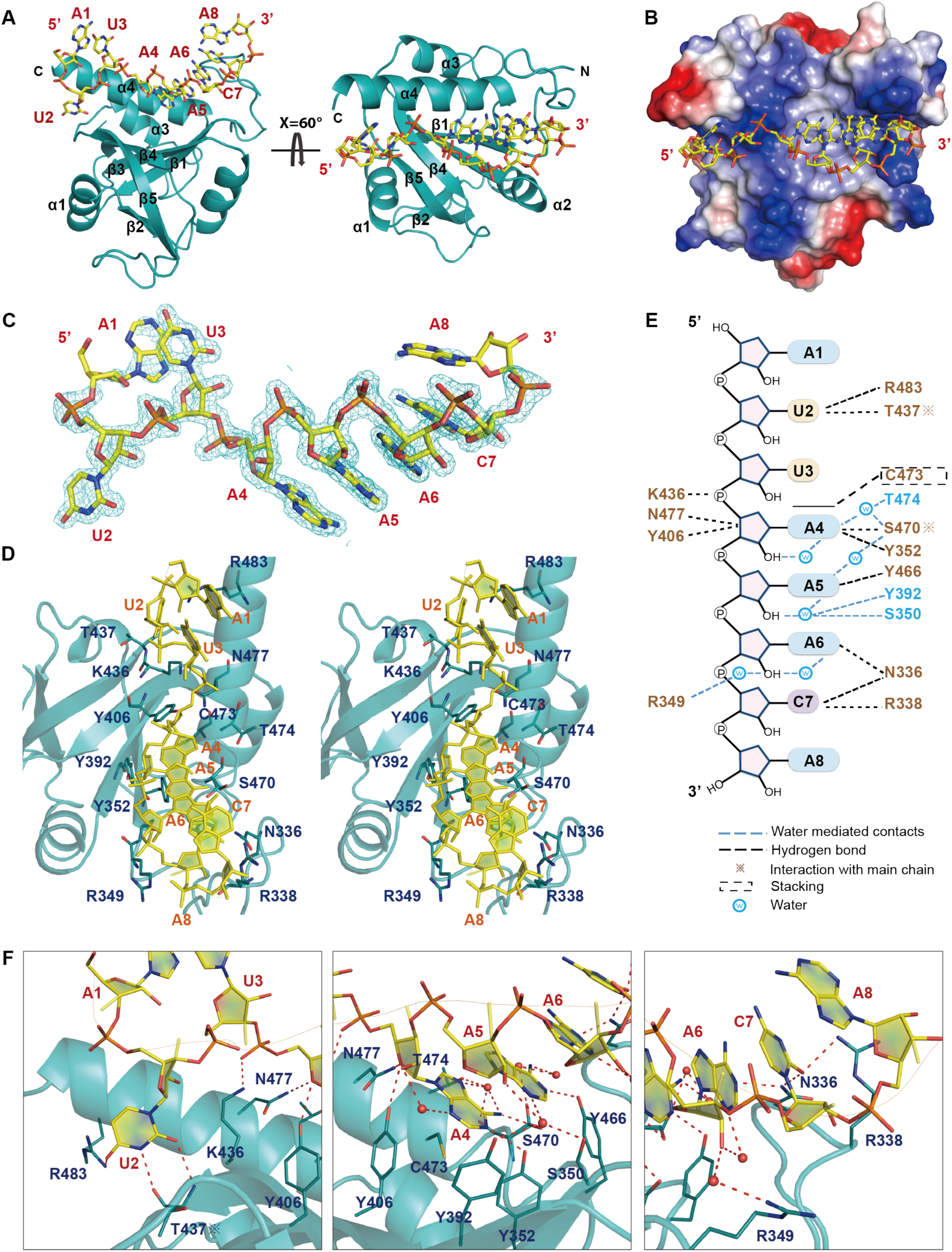
(A) Overall structure of the Mmi1 YTH domain in complex with RNA 5’-AUUAAACA-3’. The protein is shown in the cyan cartoon. The 8-mer RNA is shown in the yellow stick.[G1][G2] (B) Electrostatic surface of the Mmi1-YTH domain in the region of RNA binding. The RNA is located in a positively charged surface (blue) in the protein. The surface charge at neutral pH is displayed as blue for positive, red for negative, and white for neutral[G3]. (C) Omit F_O_-F_C_ electron density for the RNA at 1.6 Å resolution, contoured at 1.0σ. (D) Stereo view highlighting intermolecular contacts in YTH-RNA complex. The protein (in cartoon representation) is colored in cyan and RNA (in cartoon and stick representation) is colored in yellow. Residues involved in recognizing the RNA are shown in cyan sticks. (E) Schematic representation of protein-RNA interactions in the complex. Hydrogen bonds and water-mediating hydrogen bonds interactions between RNA bases and sugar-phosphate backbone with amino acid residues of YTH domain are shown by black and blue dashed lines, respectively. Snowflake symbol denotes interactions involving proteins [G4][G5] main chain atoms. (F) The close-up view of detail interactions between all nucleotides and the interacting residues of YTH domain, water molecules are shown as red spheres. Hydrogen-bonding interactions are indicated by dashed lines in red.

### Specific recognition of DSR core motif by Mmi1 YTH domain

The DSR core motif starts from U2 to C7. The uracil at the 2 position (U2) forms two base-specific hydrogen bonds with the side chain of T437 via NH_2_ group (N^3^) and carbonyl oxygen (Figures 3E and 4A). Substituting U2 for other nucleotides (cytosine) will disrupt both hydrogen bonds (Figure 4H). The uridine at the 3 position (U3) has no interaction with the Mmi1 YTH domain, consistent with a lack of selectivity at this position. The adenine at the 4 position (A4) forms three hydrogen bonds with the side chains of Y352, K436, N477, respectively (Figures 3E, 4B and 4C). The phosphate group of A4 forms a hydrogen bond with the amino group of K436, and the sugar ring forms a hydrogen bond with the amino group of N477(Figure 3E). The hydroxyl of Y352 has interaction with the NH_2_ at the N^1^ and N^6^ position of A4’s base (Figure 4B and 4C). The adenine at 5 position (A5) forms a hydrogen bond with the hydroxyl group of Y466 by NH_2_ at N^1^ position (Figure 3E and 4D), and the pentose group of A6 has an interaction with the side chain of R349 (Figure 3E and 4E). The three adenine base stacking is very special, if we replace any of them with a guanine, the binding affinities will be dramatically reduced as shown by our ITC data (Figure 4H). The cytosine at the 7 position (C7) forms a hydrogen bond with the side chain amino group of N336 and R338 by its base carbonyl oxygen (Figures 3E and 4F). The importance of the interactions has also been verified by our mutagenesis studies. Mutating R349, K436 or T437 to Alanine diminishes binding affinity shown by ITC data (Figure 4G and 4I). Interestingly, none of the residues take part in the binding of DSR core motif is conserved in the YTH domain family (Figure 1B).

**Figure 4.**
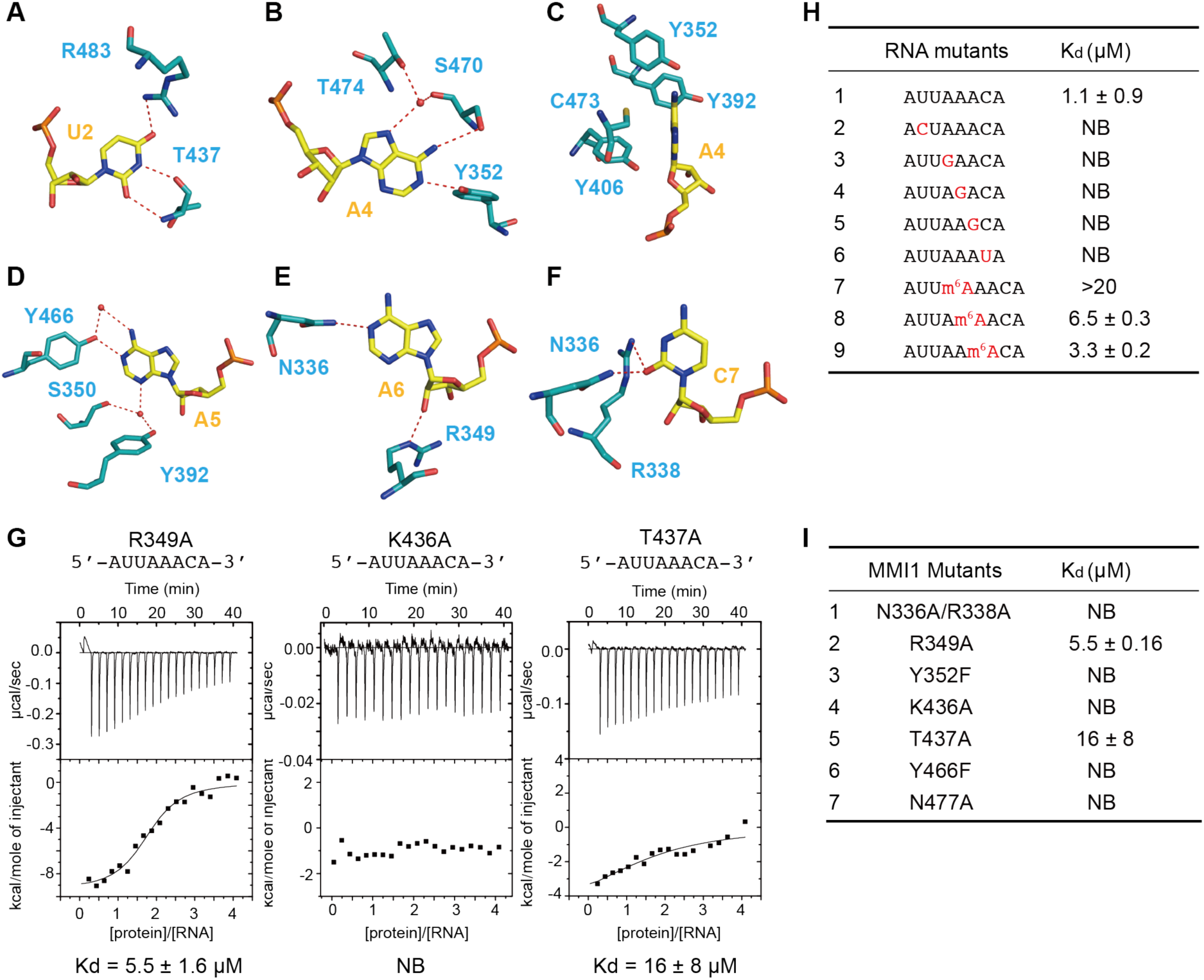
Detail intermolecular protein-RNA recognition contacts in the complex. (A) Hydrogen bonds of the O^4^ with the side chain of R483 and the O^2^ and N^3^ of U2 with the backbone of T437. (B) Hydrogen-bonding of the N^1^ and N^6^ of the A4 with the side chain of Y352 and S470. A4 N^7^ is hydrogen bonded to the T474 mediated by a water molecule. (D) N^1^ of A5 is hydrogen-bonded to the side chain of Y466. N^3^ IS hydrogen bonded to the S350 and Y392 mediated by a water molecule. (E) Hydrogen-bonding of the N^1^ and 2’-OH of the A6 with the side chain of N336 and R349. (F) Hydrogen-bonding of O^2^ of the C7 with the side chain of N336 and R338. (F) The base of A4 is surrounded by a hydrophobic environment, which is composed with C473, Y406, Y352, Y392. (G) ITC result of R349A, K436A, T437A mutants with the 8-mer RNA target. Solid lines indicate nonlinear least-squares fit the titration curve, with *ΔH* (binding enthalpy kcal mol-^1^), Ka (association constant) and *N* (number of binding sites per monomer) as variable parameters. Calculation values for K_d_ (dissociation constant) are indicated. (H) The sequence of RNA used for ITC titration study. Each nucleotide point mutant is substitute with similar purine or pyrimidine colored in red except U3. (I) Mutants of YTH domain used for Isothermal titration calorimetry study. Target RNA 5’-AUUAAACA-3’ is used.

## DISCUSSION

Meiosis is a very important stage in individual life cycle coupled with many dramatic changes, as a key factor in regulating meiotic-specific mRNAs in the mitotic stage, Mmi1 DSR selective removal system plays an essential role in fission yeast to control the expression in the correct time and space. Our structure not only illuminates the molecular basis how and why Mmi1 YTH domain binds to the DSR core motif, but also propose a new YTH-RNA recognition mechanism distinguish from the m^6^A reader so far. Here the specific YTH domain may hint us an entirely different system to carry out the same function in *S. pombe*.

## MATERIALS AND METHODS

### Protein and RNA preparation

We cloned the PCR-amplified cDNA fragments encoding fission yeast *S. pombe* Mmi1 (349–477) and Mmi1(326–488) into a modified pET28a vector that adds a Ulp1 protease-cleavage His_6_-Tag at N-terminus. The recombinant proteins were expressed in *Escherichia. Coli* strain BL21(DE3) in LB medium at 18 °C in the presence of 2 mM IPTG. The overnight cell cultures were harvested by centrifugation and dissolved in the lysis buffer containing 20 mM Tris-HCl, pH 8.0, 500 mM NaCl, 25 mM Imidazole. The cells were lysed by high pressure. The supernatant was collected after centrifugation at 17,000 rpm for 1 hour and then applied to Ni-NTA (GE). The target protein was washed with lysis buffer and then eluted with a buffer containing 20 mM Tris-HCl, pH 8.0, 500 mM NaCl and 500 mM imidazole. Ulp1 protease was added to remove the N-terminal tag and Sumo of the recombinant protein and dialyzed with lysis buffer 3 hours. The mixture was applied to another Ni-NTA resin to remove the protease and uncleaved proteins. Then the cleaved recombinant proteins were further purified by Superdex75 gel filtration (GE Healthcare) in a buffer containing 10 mM Tris-HCl, pH 8.0, 100 mM NaCl. The mutants of Mmi1 YTH domain were cloned using the overlap PCR and were expressed and purified in the same way as the wild-type. RNA oligonucleotides were commercial synthesized (Thermo Fisher Scientific, Inc.) and deprotected or synthesized by ABI 394 DNA/RNA synthesizer in our lab.

### Crystallization and data collection

For crystallization of the Mmi1 YTH domain (residues 349–479), 1 μl protein (10 mg/ml) was mixed with 1μl crystallization buffer using the hanging drop vapor diffusion method at 20 °C. The diffracting crystal was crystallized in a buffer containing 0.1 M Bis-Tris, pH 5.5, 25% PEG 3350. For crystallization of the Mmi1(326–488)-AUUAAACA complex, 10 mg/ml protein (final concentration) was mixed with the 8-mer RNA AUUAAACA (Thermo Fisher Scientific, Inc.) in a molecular ratio of 1:1.2. The mixture was incubated at room temperature for 0.5 h before crystallization. The RNA complex was crystallized under the 20%PEG3350, 8% Tacimate pH5.0. For data collection, crystals were flash frozen in the above reservoir solutions supplemented with 20% glycerol. Diffraction data were collected using a copper rotating anode X-ray source, integrated and scaled with HKL2000. SAD data were collected on Mmi1 (349–477) Se-Met crystals (2.0 Å resolution) at the wavelength 0.97988 Å, corresponding the Se absorbance edge. A native (1.49 Å resolution) dataset was collected on crystals of Mmi1 (349–477) at the wavelength 0.9795 Å, another native dataset was collected on crystals of Mmi1 (326–488) at the same wavelength. All diffraction data were collected on BL-17 U beamline. The data were processed by HKL2000. The crystals of Mmi1 (349–477) belong to space group P2_1_2_1_2_1_, and the Se-Mmi1 (349–477) crystals belong to space group P2_1_. The crystals of Mmi1 (326–488) belong to space group C222_1_ and the crystals of Mmi1(326–488)-RNA complex belong to space group P2_1_2_1_2. All of them have two molecules or protein-RNA complex per asymmetry unit.

### Structure determination and refinement

The SHELXD was used to locate the selenium sites in the crystallographic symmetry units of the Mmi1 (349–477) crystals. The program SHELXE(Sheldrick 2008) was used to calculate SAD phases. The electron-density map was calculated in the CCP4 suite. Automatic protein model building was performed with PHENIX(Adams et al. 2010). The structure of Mmi1-RNA complex was determined by molecular replacement with PHASER MR(McCoy et al. 2007) in the CCP4(Winn et al. 2011) suit using the free protein as the search model. The final models were iteratively rebuilt, refined and validated with COOT(Emsley and Cowtan 2004) and REFMAC5 (Murshudov et al. 2011) respectively. A summary of crystallographic statistics is shown in Table 1. An m|*F*o| – D|*F*c| map was calculated within the CCP4 program suite.

### ITC measurements

All ITC measurements were recorded at 25 °C using a MicroCal ITC200 (GE Healthcare). All RNAs used for ITC binding experiments were synthesized with IDT 394 DNA/RNA synthesizer. All proteins and RNAs are dialyzed or dissolved in the same buffer containing 20 mM KCl, 80 mM NaCl, 2 mM EDTA, 50 mM Tris–HCl (pH 7.5), 0.05% NP-40, 1 mM MgCl2, 2 mM dithiothreitol, 20 injections were performed by injecting 2μl 100μM proteins into a sample well containing 5 μM of RNA. The concentration of the proteins and RNAs were estimated with absorbance spectroscopy using the extinction coefficients OD280nm and OD260nm, respectively. Binding isotherms were plotted, analyzed and fitted in a one-site binding model by Origin Software (MicroCal Inc.) after subtraction of respective controls. The K_d_, entropy, enthalpy and Gibbs free energy, as well as their deviations, were also calculated using Origin Software (MicroCal Inc.) during the ITC curve fitting in a one-site model.

## DATA DEPOSITION

Coordinates have been deposited in the Protein Data Bank with accession codes for 5EIM for YTH-RNA complex and 5EIP for apo-YTH domain structures.

## SUPPLEMENTARY MATERIAL

Supplementary material is available for this article.

## ACKNOWLEDGMENTS

We thank the staff of the Beamline BL17U at SSRF for assistance with data collection. This work was supported by the National Basic Research Program of China (2011CB966304 and 2012CB910502), and by the National Natural Science Foundation of China (31230041).

